# Efficacy of RMC-6236 (Daraxonrasib) and novel combination strategies targeting resistance in RAS pathway-driven neuroblastoma

**DOI:** 10.64898/2026.06.24.734371

**Authors:** Ivette Valencia-Sama, Lynn Kee, Alex Weiss, Madeline N. Hayes, Michael Ohh, Meredith S. Irwin

**Affiliations:** Cell & Systems Biology Program, The Hospital for Sick Children, Toronto, Canada; Developmental, Stem Cell and Cancer Biology Program, The Hospital for Sick Children, Toronto, ON, Canada; Department of Molecular Genetics, University of Toronto, Toronto, ON, Canada; Department of Laboratory Medicine and Pathobiology, University of Toronto, Toronto, Canada; Department of Medical Biophysics, University of Toronto, Toronto, Canada; Department of Paediatrics, The Hospital for Sick Children, Toronto, Canada

**Keywords:** Neuroblastoma, RAS, MAPK, RMC-6236, daraxonrasib, avutometinib

## Abstract

Metastatic neuroblastoma (NB), the most common pediatric extra-cranial solid tumor, has a cure rate of <50%. DNA-sequencing studies have demonstrated rare recurrent driver mutations at diagnosis, with the most common alterations detected in ALK-RAS-MAPK pathway. Activating ALK and RAS-MAPK mutations are associated with inferior outcome and are increased at relapse, and thus, represent therapeutic vulnerabilities in NB. Previously, we identified combinations of RAS/MAPK inhibitors, including SHP2 and MEK, with efficacy in resistant MAPK-altered tumor cells, including those with the most common NB-associated *RAS* mutation NRAS-Q61K. However, toxicities of SHP2 inhibitors and promising results using compounds that directly target RAS suggest there may be superior strategies to target RAS/MAPK pathway in NB. Here, we have assessed the efficacy of RAS/MAPK inhibitors, including tovorafenib (pan-RAF), RMC-6236/daraxonrasib (pan-active-RAS) and avutometinib (RAF/MEK) in NB *in vitro* and *in vivo* using NB models harboring differing genomic status of RAS/MAPK pathway effectors. We demonstrate selective efficacy of RMC-6236 and avutometinib via RAS-MAPK pathway inhibition in NB cells and xenografts harboring *RAS, NF1* or *ALK* alterations. Importantly, we demonstrate that presence of the NRAS-Q61K mutation confers drug sensitivity. Using newly generated and previously established NB cell models of acquired resistance to RMC-6236 or the ALK inhibitor lorlatinib, we identified targeted combinations, including RMC-6236 plus avutometinib, that demonstrate re-sensitization in resistant NB cell and xenograft models. Finally, transcriptomic studies of RMC-6236-resistant cells detected upregulation of RAS/MAPK signatures, as well as TNFα/NFκB and IL-6/JAK/STAT3 pathway enrichment, thus informing future combinations to enhance sensitivity to RAS inhibitors.

**STATEMENT OF SIGNIFICANCE:** Our work demonstrates that newly available RAS pathway inhibitors RMC-6236/daraxonrasib and avutometinib have efficacy in neuroblastoma tumors, which have frequent alterations in the RAS/MAPK pathway. These drugs with early efficacy results in adult RAS-driven tumors provide an important option for patients with relapsed neuroblastoma alone or in combination.

## INTRODUCTION

Neuroblastoma (NB) is the 3rd most common pediatric cancer, but disproportionately responsible for 15% of pediatric cancer-associated mortality. Most patients with low and intermediate risk disease are cured with minimal clinical intervention; however, survival for high-risk metastatic NB patients remains under 50%, even with intensive multimodal therapeutic interventions (1–6). For patients who recur, less than 5% are cured (7, 8) and thus, new therapies are urgently needed. The main obstacles to opening Phase I and II trials for pediatric tumors, including relapsed NB, are the lack of novel therapies with efficacy in cellular and animal models that are also safe for children and available from pharmaceutical companies for pediatric clinical trials (1, 9). Thus, identifying agents developed for adult-onset cancers harboring similar genomic alterations can be an advantage to shorten the timelines for pediatric patient access.

NB is a heterogeneous tumor with rare recurrent driver mutations. Amplification of the *MYCN* oncogene, detected in 20% of tumors, predicts poor prognosis and often impacts sensitivity to anti-cancer drugs; however, thus far, there are no drugs that successfully target MYCN (10–12). The most commonly mutated gene in NB is *ALK*, which is detected in ∼10% of tumors from newly diagnosed NB (13–15). Next generation sequencing (NGS) studies have also detected less common alterations (<5%) in *MYC*, *PTPN11*, *NF1*, *H/K/NRAS, ARID1a*, and *ATRX* (16–19). Germline mutations in ALK are detected in ∼1% of NB patients (20–23) and NB tumors are detected in patients with other cancer predisposition syndromes including Neurofibromatosis and ‘RASopathies’ (e.g. Costello, Noonan), which have germline alterations in RAS pathway genes including *NF1*, *RAS* and *PTPN11* (also known as SHP2) (24–29).

Mutations in RAS/MAPK pathway genes or *ALK* (which can activate RAS/MAPK signaling) are associated with poor outcome at diagnosis (14, 30–32). Notably, NGS studies have shown that ALK and RAS/MAPK alterations (including *RAS, PTPN11, NF1*) are enriched in tumors and ctDNA obtained at the time of relapse. (33–38). While *NRAS* mutations (specifically Q61K) are the most common, other alterations such as *K/HRAS*, *NF1*, and *PTPN11* mutations have been detected in NB tumors (17–19, 39) and ctDNA (30, 37, 38). These results suggest that pharmacologic targeting of ALK/RAS/MAPK pathway may be beneficial for the treatment of recurrent NB; however, to date, pediatric clinical trials are only available to target *ALK* mutations (1, 40–42).

*RAS* oncogenes *KRAS*, *HRAS* and *NRAS* encode membrane-bound GTPases that cycle between GTP-bound active and GDP-bound inactive states to regulate activation of downstream effectors RAF (ARAF/BRAF/CRAF), MEK and ERK, and subsequently modulate cellular proliferation, differentiation and survival (43). Upstream positive regulators include the *PTPN11*-encoded tyrosine phosphatase SHP2, which increases RAS-RAF binding via RAS dephosphorylation and activates the RAS/MAPK pathway (43, 44). In cancer, the RAS/MAPK pathway is commonly hyperactivated as a result from gain-of-function (GOF) mutations in upstream tyrosine kinase receptors (RTKs) including *ALK*; loss-of-function (LOF) mutations of negative regulators such as *NF1*; or activating *RAS* mutations (e.g. hotspot codons 12, 13 and 61) which are detected in ∼30% of adult-onset cancers (45–48).

Therapeutic strategies targeting activated RAS/MAPK signaling include direct RAS inhibition (49–53), vertical MAPK pathway strategies (54–58), and synergistic combinations with alternate pathway inhibitors (e.g. PARP, CDK4/6, RTKs) (59–66). Previously, we and others have investigated the efficacy of single agent and combinations targeting various effectors of RAS signaling, including combinations of various SHP2 inhibitors with ALK, MEK or ERK inhibitors (67, 68). However, SHP2i monotherapies have demonstrated toxicity in Phase I/II adult trials, potentially impacting further development. Furthermore, promising pre-clinical and early phase data in adult trials have demonstrated efficacy for novel RAS-targeting agents, such as the tri-complex RAS-ON inhibitor RMC-6236 (daraxonrasib). RMC-6236 works by forming a tri-complex with the intracellular chaperon CYPA and RAS-GTP, which blocks RAS effector binding and halts downstream MAPK signaling (69). Importantly, since it inhibits active RAS proteins regardless of isoform or mutational status, RMC-6236 has shown robust anti-tumor activity in preclinical models of *KRAS*-mutant non-small cell lung cancer (NSCLC), colorectal cancer (CRC), and pancreatic ductal adenocarcinoma (PDAC) including metastatic *RAS*-mutant PDAC (70, 71). Ongoing clinical trials of RMC-6236 alone or in combination with chemotherapy have thus far demonstrated promising efficacy and tolerability in patients with advanced or metastatic NSCLC, PDAC, CRC and melanoma tumors harboring *N/H/KRAS* mutations (NCT05379985, NCT06162221, NCT06625320, NCT06881784, NCT07252232), and has led to a recent FDA-authorized expansion access approval for patients with previously treated metastatic PDAC (NCT07573215).

Other agents targeting downstream MAPK components have shown promising results in adult and pediatric tumors. Tovorafenib, a selective CNS-penetrant pan-RAF type II inhibitor, has shown single-agent efficacy in melanoma, spindle cell sarcoma and pediatric low-grade glioma (pLGG) tumors harboring *RAF* fusions or mutations (72–76), and demonstrated synergistic activity with anti-MEK therapy in *NF1*-altered rhabdomyosarcomas and malignant peripheral nerve sheath tumors (73). Currently, tovorafenib has been granted FDA approval for the treatment of *BRAF*-altered pLGG with ongoing trials for various MAPK-altered solid tumors (NCT05760586, NCT07206849, NCT04985604, NCT04775485, NCT07121829).

The anti-MAPK agent avutometinib is a first-in-class RAS/MEK clamp inhibitor that simultaneously blocks MEK catalytic activity as well as RAF-induced phosphorylation by promoting the formation of inactive RAF-MEK complexes (77, 78). Preclinical and clinical efficacy has been reported in MAPK-altered low grade serous ovarian cancer, uterine carcinosarcomas, high-grade endometrioid endometrial cancer, and multiple myeloma, as well as *KRAS*-driven PDAC and NSCLC tumors (79–84). Recently, avutometinib has been granted FDA-approval in combination with the FAK inhibitor defactinib for the treatment of patients with ovarian cancer (NCT06072781) (85, 86), and over 20 clinical trials are currently evaluating avutometinib as a monotherapy or in combination regimens in other MAPK-driven solid tumors including neuroblastoma (NCT04620330, NCT06104488).

Here we report single agent and combination efficacy for these agents and demonstrate that sensitivity is determined by mutational status of *RAS* or MAPK effectors. RMC-6236 and ALK inhibitor resistant cell models were generated and used to demonstrate re-sensitization with combinations including RMC-6236 with avutometinib *in vitro* and *in vivo*. Finally, RAS/MAPK, TNFα/NFκB and IL-6/JAK/STAT3 gene signatures were enriched in RMC-6236 resistant cells, potentially supporting identification of additional novel combination strategies for therapy-resistant NB. These findings suggest that RAS-targeting agent strategies should be prioritized for further development, especially in patients with alterations in the RAS/MAPK pathway.

## MATERIALS AND METHODS

### Cell lines

Human neuroblastoma cell lines SK-N-SH, LAN-6, SK-N-FI, Kelly, SH-EP, IMR-32, SK-N-BE, SK-N-DZ, CHP-212, SK-N-AS, COG-N-415 and COG-N-519 were purchased from ATCC or obtained from COG Childhood Cancer Repository (http://cccells.org). NB-EB cell line was provided by Dr. Marielle E. Yohe (National Cancer Institute, NIH). Isogenic SH-EP stable cell lines were engineered to overexpress pcDNA3-NRAS-WT-3xflag (SH-EP-NRAS^WT^), pcDNA3-NRAS-Q61K-3xflag (SH-EP-NRAS^Q61K^), or an empty pcDNA3 vector (SH-EP-EV) as described (67). Short tandem repeat (STR) cell authentication (TCAG Sequencing Facility, Toronto) and *Mycoplasma* testing (InvivoGen) was performed prior to experimental use.

### Cell culture

IMR-32 and CHP-212 cells were cultured in EMEM media (Wisent); SK-N-AS, SK-N-DZ and SK-N-F1 cells were cultured in DMEM (Wisent); SK-N-SH, LAN-6, Kelly, SH-EP, SK-N-BE and NB-EB were cultured in RPMI media (Wisent). All media was supplemented with 10% FBS, 3mM L-glutamine (Gibco), 0.1mM nonessential amino acids (Gibco), and 1mM sodium pyruvate (Gibco). COG-N-415 and COG-N-519 cells were cultured in IMDM (Wisent) supplemented with 20% FBS (Wisent), 3mM L-glutamine (Gibco), and 1x ITS (5 μg/mL insulin, 5 μg/mL transferrin, 5 ng/mL selenous acid). Cells were incubated at 37°C and 5% CO2 tissue culture incubators.

### Chemicals

TNO155 (#HY-136173), Tovorafenib (#HY-15246), RMC-6236/daraxonrasib (#HY-148439), Avutometinib (#HY-18652), Olaparib (#HY-10162) and Ribociclib (#HY-15777) were purchased from MedChemExpress. Inhibitors were resuspended in DMSO according to manufacturer’s instructions. For murine experiments, RMC-6236/daraxonrasib (#HY-148439) and Avutometinib (#HY-18652) were resuspended in final concentration of 10% DMSO + 20% PEG400 + 10% Solutol HS15 in water.

### Cell viability assays

A total of 2.5-3 x 10^3^ cells per well were treated with inhibitors in 96-well plates for 120 hours. PrestoBlue reagent (Invitrogen) was added for 2 hours, and fluorescence was assessed using a microplate reader (BioTek Synergy LX) with a l530 excitation/l590 emission filter. IC_50_ curves were calculated using GraphPad Prism v.11.0.2 (GraphPad Software, LLC). For each condition, experiments included 3-6 technical replicates and were performed ≥ 3 times.

### RAS-GTP pulldowns and immunoblotting

Cells were lysed in RIPA buffer (50 mmol/L Tris, pH 8.0, 150 mmol/L NaCl, 1% NP-40, 0.5% deoxycholate, 0.1% SDS) or EBC buffer (50 mmol/L Tris, pH 8.0, 120 mmol/L NaCl, 0.5% NP-40) and supplemented with EDTA-free protease inhibitor cocktail and PhosSTOP phosphatase inhibitor cocktail. For RAS-GTP pulldowns, cells were lysed in EBC buffer, quantified, and ≥1 mg of whole cell extraction (WCE) lysates were precipitated using immobilized GTP-beads (Jena Bioscience) and incubated at 4°C with overnight rocking as previously described (67). Bound proteins were washed five times in NETN buffer (20 mmol/L Tris, pH 8.0, 100 mmol/L NaCl, 1 mmol/L EDTA, 0.5% NP-40) and eluted by boiling in 1x sample buffer. For immunoblotting analyses, cells were lysed in RIPA buffer, quantified, resolved, electro-transferred and probed with the indicated antibodies as previously described (67).

### Antibodies

Rabbit monoclonal or polyclonal antibodies against p-ERK1/2 (T202/Y204) (#9101, 1:1000), p-MEK1/2 (S217/221) (#9121, 1:1000), MEK1/2 (#9122, 1:1000), p-BRAF (S445) (#2696, 1:1000), p-ARAF (S299) (#4431, 1:1000), p-CRAF (S289/296) (#9431, 1:1000), BRAF (#9433, 1:1000), ARAF (#4432, 1:1000), CRAF (#9422, 1:1000), NF1 (#14623, 1:1000), cleaved PARP (#9541, 1:1000) were obtained from Cell Signaling Technologies. Mouse monoclonal antibodies against pan-RAS (#OP40, 1:500), Vinculin (#05386, 1:2000) was obtained from Millipore. FLAG-M2 (#F1804, 1:1000) and β-actin (#A5316, 1:2000) were obtained from Sigma. ERK1/2 (#9107, 1:1000) was obtained from Cell Signaling Technologies.

### Statistical analyses

Unpaired two-tailed Student t-test was used to assess statistical significance between two treatment groups. Analysis of variance (ANOVA) followed by a post-hoc Tukey test was used for multiple group comparisons. For Kaplan-Meier survival curves, log-rank (Mantel-Cox) tests were performed. Data reported as mean and standard deviation (SD) represents at least two independently conducted experiments. Statistical analyses were performed using GraphPad Prism v.11.0.2 (GraphPad Software, LLC), and p-values < 0.05 were considered significant.

### Animal studies

Female NOD/SCID mice (6–8 weeks) were injected subcutaneously with 1 x 10^6^ SK-N-AS or SK-N-AS-RR (RMC-6236 resistant) cells in 0.1 mL suspension containing 50% of Geltrex (Gibco) or Cultrex (R&D) in PBS. Once tumors reached ∼100 mm^3^ mice were randomized to receive a 0.2 mL suspension containing either vehicle or RMC-6236 (25 mg/kg once per day, q.d.) alone; or avutometinib (0.3 mg/kg q.d. or every other day, q.o.d.) alone or in combination with RMC-6236 by oral gavage 5 days per week, similar to other treatment regimens (69, 71, 81, 83, 87). Tumor volumes were monitored at least twice per week, and endpoint was defined as measurements over 10 mm in two of three volume dimensions. All animal studies were performed in accordance with University Health Network and SickKids Institutional Animal Utilization Protocol guidelines.

### Bulk RNA sequencing and processing

After sequencing, FastQC(0.12.1) was used to confirm quality and FastP(0.24.1) for adapter trimming/processing with arguments length_required 20 and qualified_quality_phred 15 supplied. A decoy-aware Salmon index was prepared using release 114 of the human transcriptome and genome from ensembl with –k 23 argument supplied. After index preparation, counts were generated with Salmon’s quant function and the following arguements: validateMappings, rangeFactorizationBins 4, gcBias, seqBias. Quantifications were processed in R. Briefly, tximeta(1.27.1) was used to prepare a SummarizedExperiment object which was then processed with summarizeToGene and subsetted for each cell line for use with DESeq2(1.48.2). Genes with less than 10 counts across replicates were dropped. Ashr(2.2) was used to scale log2foldchange values based on standard error, and these values were then used to create a ranked list for GSEA. Pheatmap(1.10.13) was used to highlight the scaled (using scale = “row”) expression of leading edge genes from vst() transformed counts. For human cell lines, GSEAs desktop application(4.4.0) was used to score sets from MSigDB, as well as custom gene sets taken from existing literature.

### Data availability

All data generated are available upon request from the corresponding author.

## RESULTS

### Selective efficacy of RMC-6236 and avutometinib in RAS/MAPK-altered neuroblastoma cells

A panel of thirteen neuroblastoma cell lines with various MAPK pathway alterations, including *RAS* and *ALK*, were selected to assess sensitivity to inhibitors that directly target RAS or MAPK pathway components and are currently approved or in early phase human trials (**Table 1**). Inhibitors included the pan-RAF (type II) inhibitor tovorafenib, the GTP-bound RAS(ON) inhibitor RMC-6236, the RAF/MEK clamp inhibitor avutometinib, and the SHP2 inhibitor TNO155. As we observed previously (68), cells harboring *ALK*, *NF1* or *KRAS-G12X* alterations were sensitive to the SHP2i TNO155, whereas *NRAS-Q61K* status correlated with SHP2i resistance (**Fig. 1A**, **Table 1**). Sensitivity to the FDA-approved tovorafenib, which has demonstrated efficacy in pediatric glioma patients with *NRAS* mutations (76, 88), was similar across most NB cell lines tested, regardless of RAS/MAPK status (**Fig. 1B**, **Table 1**). In contrast, both RMC-6236 and avutometinib showed increased efficacy in cells harboring *RAS* mutations (including *NRAS-Q61K*) or *NF1* alterations, and in most cells bearing *ALK* mutations, compared to the least sensitive lines with no Ras-associated mutations (no-Rm) (**Fig. 1C** and **1D**, **Table 1**). Notably, most of these effective concentrations were similar to those published for adult tumor models (69, 71). These data, particularly those with RMC-6236, uncover novel candidates to test in larger panels of NB cells with RAS activation (e.g. *RAS* and *NF1* genomic alterations).

**Figure 1.**
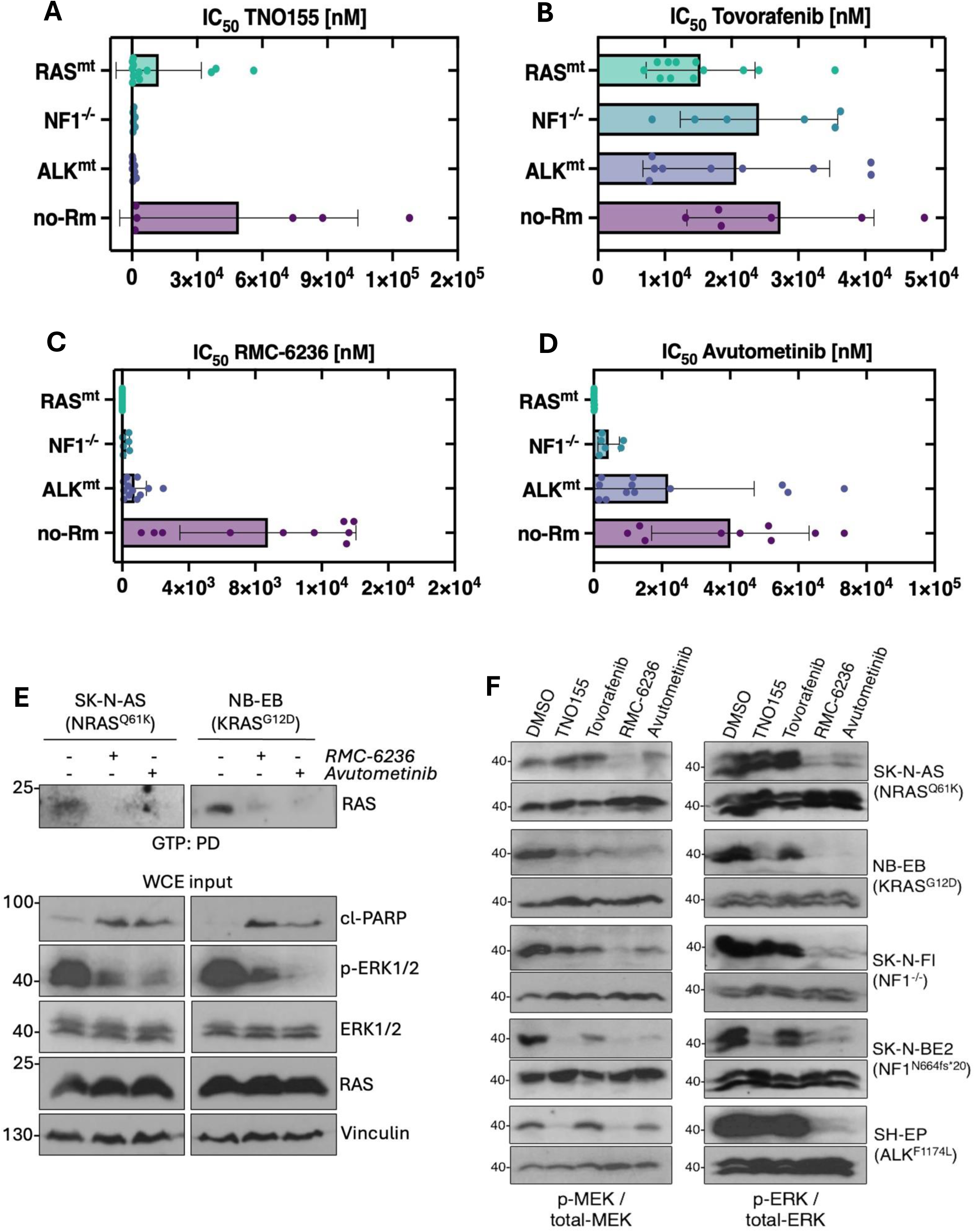
RAS/MAPK-altered NB cells are sensitive to RAS pathways inhibitors. **A**-**D**, Calculation of IC_50_ (n=3) in NB cells grouped by mutational status: *RAS* mutations (RAS^mt^), *NF1* alterations (NF1^-/-^), *ALK* mutations (ALK^mt^), or no RAS-associated mutations (no-Rm), and treated with TNO155 (**A**), tovorafenib (**B**), RMC-6236 (**C**), or avutometinib (**D**) for 120 hours. **E**, GTP pulldowns (PD) of *RAS*-mutant NB cells treated with RMC-6236 [0.2 μM] or avutometinib [0.5 μM] for 24 hours. **F**, Whole-cell lysate (WCE) immunoblots of *RAS*-, *NF1*– or *ALK*-altered NB cells treated with TNO155 [1 μM], tovorafenib [2.5 μM], RMC-6236 [0.2 μM], or avutometinib [0.5 μM] for 24 hours.

**Table 1.** Mutational status and IC_50_ of NB cell lines treated with RAS/MAPK inhibitors.

| Cell line | Average IC50 [μM] (120h) |  |  |  | Mutation/Alteration |  |  |  |  |  |
| --- | --- | --- | --- | --- | --- | --- | --- | --- | --- | --- |
|  | RMC-6236 | Avutometinib | Tovorafenib | TNO155 | RAS | NF1 | MYCN | ALK | TP53 | Other |
| CHP-212 | 0.0012 | 0.0158 | 9.5613 | 4.5327 | NRAS (Q61K) | - | Amp | - | - | - |
| SK-N-AS | 0.0003 | 0.0580 | 14.3087 | 43.1520 | NRAS (Q61K) | - | - | - | H168R | - |
| NB-EB | 0.0024 | 0.1698 | 12.4053 | 0.6232 | KRAS (G12D) | - | Amp | - | - | - |
| LAN-6 | 0.0011 | 0.0724 | 25.1223 | 0.1473 | KRAS (G12C) | - | - | D1091N | P152L | p14ARF-/- |
| SK-N-FI | 0.0443 | 6.6853 | 13.9687 | 0.4392 | - | Loss | - | - | M246R | - |
| SK-N-BE(2) | 0.4094 | 2.0543 | 34.2417 | 1.1987 | - | N664fs*20 | Amp | - | C135F | - |
| SK-N-SH | 0.0813 | 8.2417 | 8.7217 | 0.1762 | - | - | - | F1174L | - | - |
| Kelly | 1.4859 | 15.2153 | 38.0147 | 0.5342 | - | - | Amp | F1174L | P177T | - |
| SH-EP | 0.4214 | 1.7973 | 15.4143 | 1.4700 | - | - | - | F1174L | - | p14ARF-/- |
| COG-N-415 | 0.9595 | 61.9003 | n/a | n/a | - | - | Amp | F1174L |  | TERT+ |
| IMR-32 | 1.8260 | 12.7950 | 16.5367 | 1.7850 | - | - | Amp | - | - | - |
| SK-N-DZ | 9.9390 | 63.2343 | 38.1170 | 92.2327 | - | - | Amp | - | R110L | CDKN2A-/- |
| COG-N-519 | 12.9850 | 44.1063 | n/a | n/a | - | - | Amp | - | - | TERT+ |

To confirm that these RAS/MAPK inhibitors suppressed downstream signaling, we asked whether they decreased activation of RAS (via RAS-GTP) and downstream MAPK targets by immune-blotting treated cells. Decreased RAS activity (GTP binding) and increased apoptosis (cleaved-PARP) were observed in *RAS*-mutant NB cells following treatment with RMC-6236 and avutometinib (**Fig. 1E**). Furthermore, both avutometinib and RMC-6236 potently reduced MAPK pathway activity (via phosphorylated ERK1/2 and MEK1/2) in NB cell lines harboring mutations in *RAS*, *NF1*, or *ALK* (**Fig. 1F**).

### NRAS-Q61K mutation confers higher sensitivity to RMC-6236 and avutometinib

Previous studies have demonstrated that NRAS-Q61K mutation is associated with resistance to some single agent RAS pathway inhibitors including SHP2i. However, since NB cells harboring Q61K mutations were sensitive to RMC-6236 and avutometinib, we examined whether NRAS-Q61K mutation specifically confers sensitivity to these inhibitors using our previously generated SH-EP isogenic cell lines overexpressing either NRAS wild-type (WT), mutant (Q61K), or an empty vector control (EV) (67). Expression of endogenous RAS or exogenous (flag-tagged) NRAS expression was confirmed (**Fig. 2A**). Concordant with previous observations in cells harboring endogenous NRAS-Q61K (67, 68, 89), SHEP-Q61K cells were significantly more resistant to TNO155 compared to its WT counterpart or EV control, as determined by increased fold-IC_50_ (**Fig. 2B**). Furthermore, tovorafenib was slightly (but not significantly) more effective in SHEP-Q61K cells (**Fig. 2C**). In contrast, we observed a 3-fold and 2-fold decrease in IC_50_ (cell viability) in Q61K-expressing cells treated with RMC-6236 and avutometinib, respectively (**Fig. 2D** and **2E**). This was accompanied by increased apoptotic activity as determined by cleaved PARP expression in Q61K-overexpressing cells upon treatment with either agent (**Fig. 2F**). These data support RMC-6236 and avutometinib as promising agents for RAS/MAPK-driven NB, including those harboring the most resistant *NRAS-Q61K* alteration.

**Figure 2.**
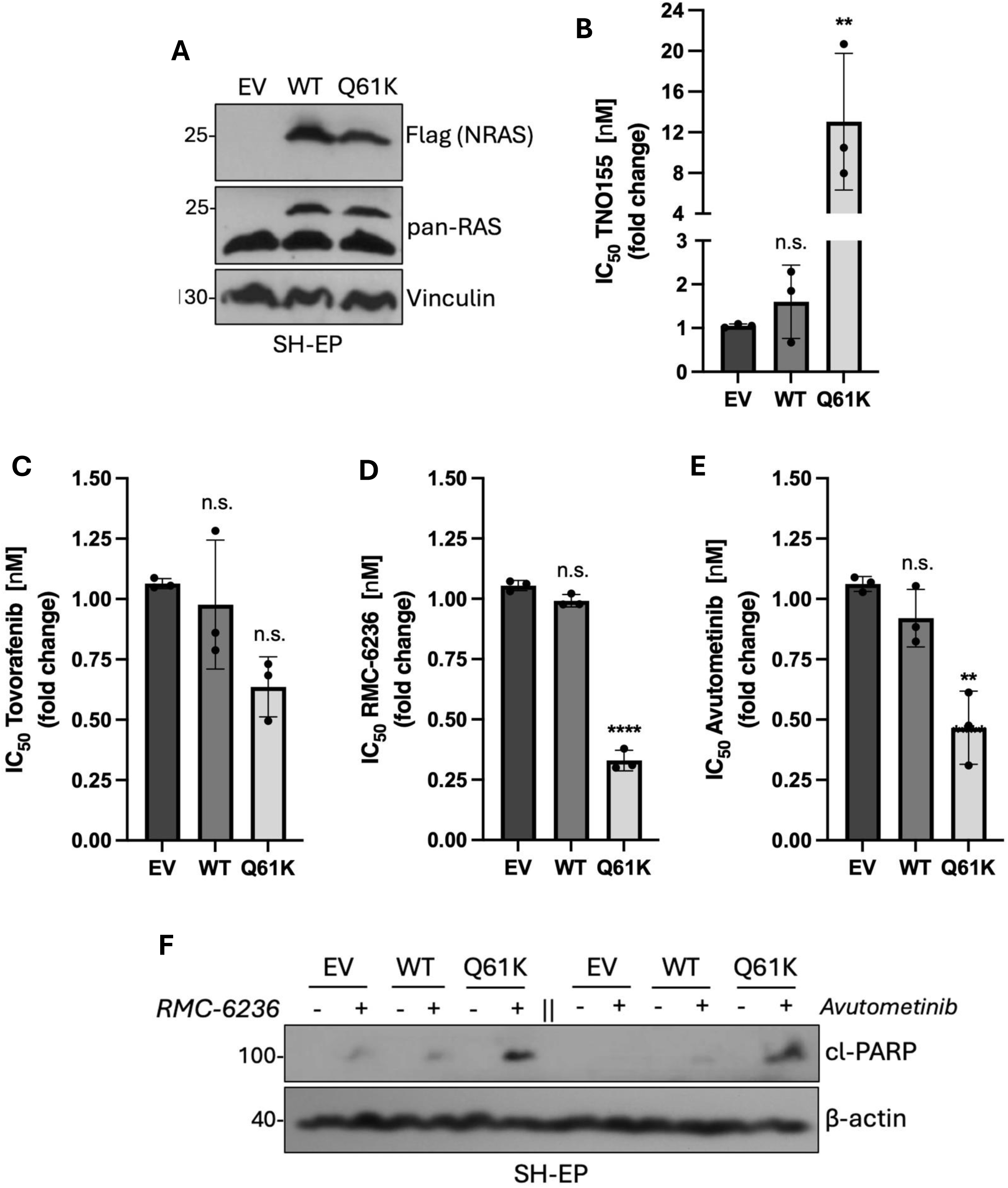
NRAS-Q61K mutation confers sensitivity to RMC-6236 and avutometinib. **A**, Isogenic SH-EP cells stably over-expressing NRAS wildtype (WT), NRAS mutant (Q61K) or an empty vector (EV). **B**-**E**, Calculation of IC_50_ fold change (relative to EV control) in SH-EP stable cells treated with TNO155 (**B**), tovorafenib (**C**), RMC-6236 (**D**), or avutometinib (**E**) for 120 hours. **F**, Immunoblots of SH-EP stable cells treated with RMC-6236 [1 μM] or avutometinib [2 μM] for 24 hours (F). **p<0.01, ****p<0.0001.

### Efficacy of RMC-6236 and subsequent acquired resistance in murine neuroblastoma xenografts

To assess RMC-6236 efficacy *in vivo*, tumor volume was monitored in mice with subcutaneous SK-N-AS xenografts harboring *NRAS-Q61K* and treated with RMC-6236 (25 mg/kg, q.d.). RMC-6236-treated xenografts demonstrated initial tumor growth reduction, followed by complete and sustained regression in 100% of tumors by day 12 (**Fig. 3A** and **3B**). In contrast, the vehicle-control group reached end point after 22 days. To evaluate the durability of RMC-6236 responses, treatment was discontinued at day 33 when tumors were non-palpable, and tumor re-growth and end-point free survival was monitored. Tumor recurrence was observed in all mice following treatment suspension (**Fig. 3B**); however, the median survival was extended by 31 days compared to control (**Fig. 3C**). Importantly, we assessed whether RMC-6236 was also effective in other NB xenografts harboring different mutations; NB-EB (*KRAS-G12D*) and SK-N-BE2 (*NF^-/-^*) xenografted tumors regressed within one week of RMC-6236 treatment compared to control (**Supplementary Fig. S1A-C**).

**Figure 3.**
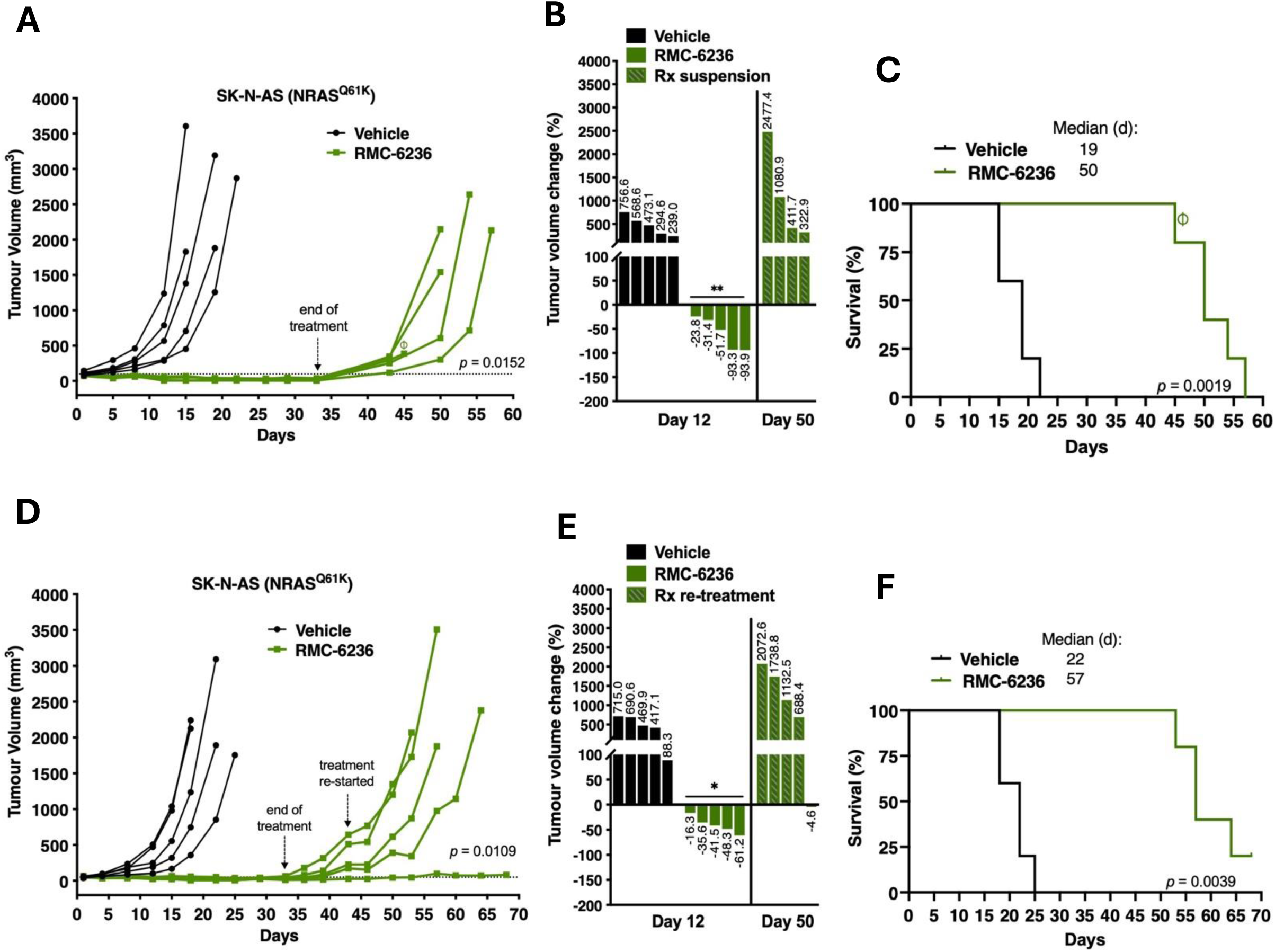
*In vivo* efficacy of RMC-6236 in RAS-mutant xenografts. **A,** Tumor volumes of SK-N-AS xenografts (NRAS^Q61K^) treated with vehicle (n=5) or RMC-6236 (25 mg/kg, q.d., n=5) and followed by treatment suspension at day 33. **B**, Percent tumor change (relative to pre-treatment) following treatment with vehicle or RMC-6236 for days 12, and after RMC-6236 (Rx) treatment suspension at day 50. **C**, Kaplan-Meier curve of endpoint-free survival. ⏀, death at day 45 was unrelated to tumor burden or drug toxicity. **D,** Tumor volumes of SK-N-AS xenografts following treatment with vehicle (n=5) or RMC-6236 (25 mg/kg, q.d., n=5); followed by treatment suspension at day 33, and re-treatment at day 43 until end-point. **E**, Percent tumor change (relative to pre-treatment) following treatment with vehicle or RMC-6236 for days 12, and after RMC-6236 (Rx) re-treatment at day 50. **F**, Kaplan-Meier curve of endpoint-free survival.

Since treatment suspension led to tumor recurrence in SK-N-AS xenografts, we examined whether sensitivity to RMC-6236 was impacted after tumor re-growth. Following a similar treatment regimen, we treated xenografts with RMC-6236 (25 mg/kg, q.d.) or vehicle for ∼5 weeks until tumors were no longer palpable, followed by a 10-day treatment suspension to allow for tumor re-growth. Once tumors recurred (day 43) treatment was re-initiated. Similar to previous observations, 100% of tumors achieved complete regression within 12 days of initial RMC-6236 treatment (**Fig. 3D** and **3E**). However, in response to re-treatment with a second round of RMC-6236, tumors were less sensitive; 4/5 of tumors showed moderate growth delay and only 1/5 tumor achieved full regression (**Fig. 3D**). Moreover after 50 days, 80% of tumor volumes were comparable to those of mice that only received one round of treatment (**Fig. 3E** vs. **Fig. 3B**). Although a second round of treatment extended the median survival by 35 days compared to control (**Fig. 3F**), the overall measured response of these re-treated tumors was less in comparisons to tumors that received only a single treatment regimen (**Fig. 3C**) These results suggest that although RMC-6236 is initially effective in *NRAS-Q61K*-bearing NB tumors, drug efficacy may be diminished after long-term use potentially due to acquired mechanisms of resistance or selection of intrinsically resistant subpopulations.

### Avutometinib combination restores sensitivity in RMC-6236-resistant neuroblastoma cells

To further examine the mechanisms of acquired resistance to RMC-6236 and identify novel treatment strategies to potentially reverse or delay resistance, we generated a RMC-6236-resistant cell line model. SK-N-AS cells were exposed to gradual dose increases (0.15 – 8.0 nM) of RMC-6236 or DMSO (vehicle) over 23 weeks, to select for RMC-6236-resistant (SKNAS-RR) or –sensitive (SKNAS-Rs) cell subpopulations (**Fig. 4A**). Acquired resistance to RMC-6236 was monitored over time and an average ∼6-fold IC_50_ increase was observed in SKNAS-RR cells compared to sensitive cells (**Fig. 4B**). Additionally, assessment of basal MAPK pathway activity showed increased activation of the downstream components ERK1/2, MEK1/2 and CRAF (also known was RAF-1) in SKNAS-RR cells compared to SKNAS-Rs cells, thus suggesting that resistance to RMC-6236 might correlate with adaptive MAPK re-activation (**Fig. 4C**).

**Figure 4.**
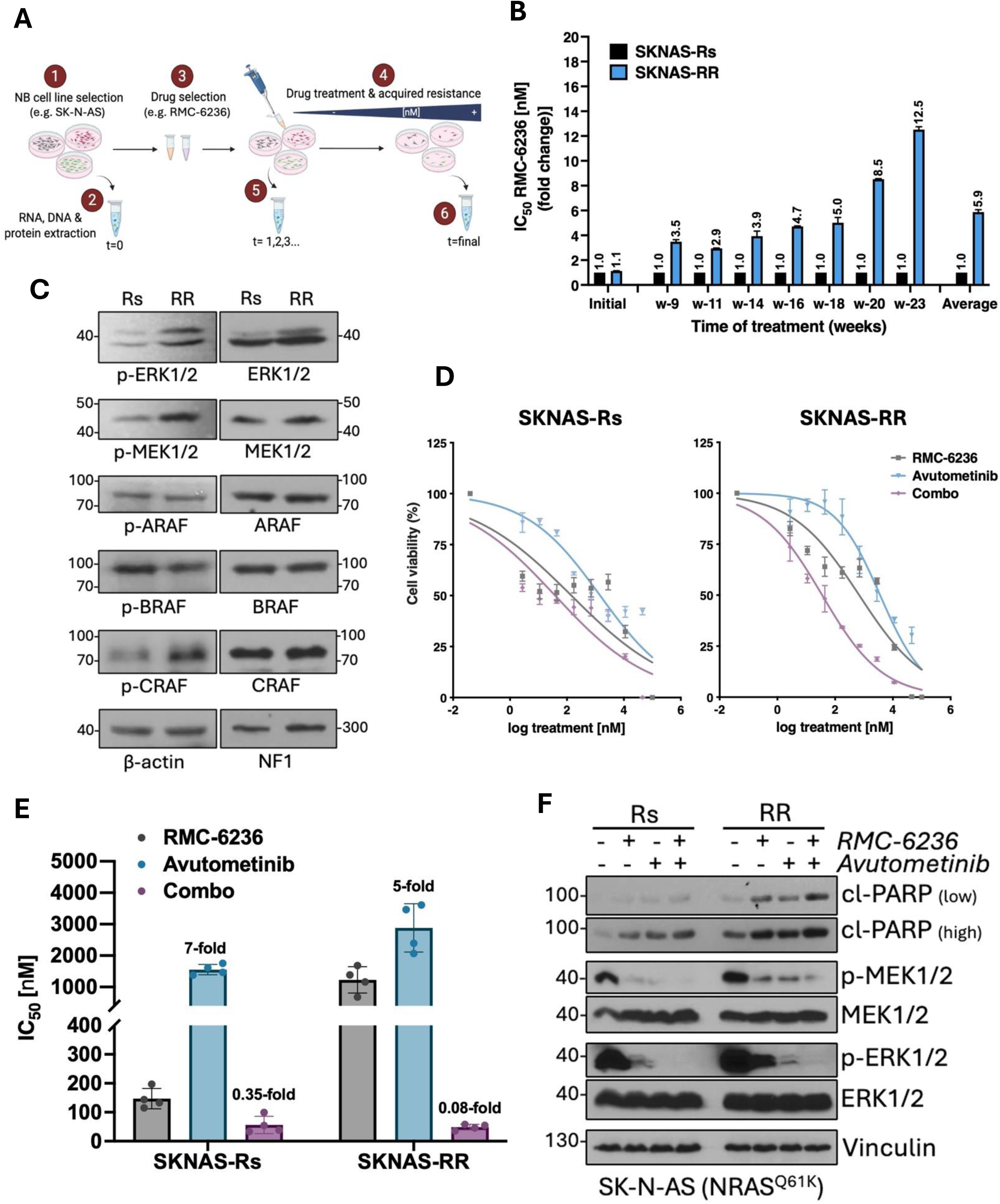
Avutometinib restores sensitivity in *RAS*-mutant cells with acquired resistance to RMC-6236. **A**, SK-N-AS cells were treated with increasing doses of RMC-6236 (0.15 – 8.0 nM) or DMSO over 23 weeks to select for subpopulations of RMC-6236-resistant (SKNAS-RR) or RMC-6236-sensitive (SKNAS-Rs) cells. Figure created with BioRender.com. **B**, Acquired resistance RMC-6236 was assessed at different time points by comparing IC_50_ fold change of SKNAS-RR cells relative to SKNAS-Rs cells (fold change). **C**, Basal expression of active (phosphorylated) or total MAPK components in SKNAS-Rs and –RR cells assessed by western immunoblots. **D**, Cell viability of SKNAS-Rs and –RR cells treated with indicated range of concentrations of RMC-6236, avutometinib or RMC-6236 plus avutometinib (Combo) for 120 hours. **E**, Average IC_50_ and fold change (relative to RMC-6236 alone) of SKNAS-Rs and –RR cells treated with RMC-6236, avutometinib or Combo for 120 hours (n=4). **F**, Immunoblots of whole cell lysates from SKNAS-Rs and SKNAS-RR cells treated with RMC-6236 [1 μM], avutometinib [2 μM] or the combination for 24 hours.

To examine whether basal sensitivity to RMC-6236 could be restored, we assessed novel combination treatments with other pathways inhibitors, including TNO155, tovorafenib or avutometinib, in SKNAS-RR cells. Addition of TNO155 showed negligible additivity effects in SKNAS-Rs cells (1.3-fold), but a mild restoration of RMC-6236 sensitivity in SKNAS-RR cells (0.4-fold) compared to RMC-6236 alone (**Supplementary Fig. S2A** and **S2B**). In contrast, tovorafenib combinations did not restore sensitivity in SKNAS-RR cells (**Supplementary Fig. S2C**). Interestingly, despite SKNAS-RR cells showing increased resistance to avutometinib alone, the combination more potently reduced cell viability and restored sensitivity to RMC-6236 (0.08-fold) as compared to SKNAS-Rs cells (0.35-fold) (**Fig. 4D** and **4E, Supplementary Fig. S2D**). In line with these observations, SKNAS-RR cells treated with the RMC-6236 plus avutometinib combination showed decreased MAPK signaling activity and increased apoptosis, compared to single agent treatments (**Fig. 4F**). These data suggest that under the pressure of treatment, acquisition of genetic alterations including *de novo* RAS/MAPK pathway mutations, could be present in SKNAS-RR cells which renders them vulnerable to avutometinib treatment combinations.

To determine whether cells with acquired resistance to other RAS/MAPK pathway inhibitors can be reversed with avutometinib, we utilized our previously established ALK inhibitor lorlatinib-resistant Kelly-LR cell model, which harbors a *de novo BRAF-G466E* mutation (68). We first assessed cell viability following TNO155, tovorafenib, RMC-6236 or avutometinib treatments alone, and observed an increased resistance to both TNO155 and RMC-6236 in Kelly-LR cells in comparison to lorlatinib-sensitive Kelly-LS cells (**Supplementary Fig. S3A**). Furthermore, neither tovorafenib nor avutometinib monotherapies showed enhanced efficacy in Kelly-LR cells compared to its sensitive Kelly-LS counterpart. Similar to SKNAS-RR cells, tovorafenib plus RMC-6236 combination showed an antagonist effect in Kelly-LR cells compared to RMC-6236 alone (**Supplementary Fig. S3B**). In contrast, the combination treatment with avutometinib potently re-sensitized Kelly-LR cells to RMC-6236 (0.04-fold) and reduced IC_50_ values to those similar to Kelly-LS cells (**Supplementary Fig. S3C**), thus suggesting that RMC-6236 resistance via MAPK-pathway mutations can be overcome by combination treatments with avutometinib. Finally, to further assess the synergistic potential of avutometinib we selected three NB cell lines (Kelly, IMR-32 and SK-N-DZ), which at baseline were less sensitive to RMC-6236 (**Table 1**), and subjected these cell lines to combination treatments with RMC-6236 and avutometinib. Interestingly, both Kelly and IMR-32 cells appeared to have improved responses using the combination treatment, as resistance to RMC-6236 was decreased by 0.13-fold and 0.41-fold, respectively (**Supplementary Fig. S4A** and **S4B**). In contrast, the combination had a negligible effect in SK-N-DZ cells (**Supplementary Fig. S4C**).

Since recent reports have suggested the use of other non-RAS/MAPK pathway targeted agents that show synergy with RAS-pathway inhibitors, including PARP and CDK4/6 inhibitors (59, 60, 66, 90), we have also assessed whether olaparib (PARPi) or ribociclib (CDK4/6i) can increase sensitivity to RMC-6236 in SKNAS-RR cells. While the olaparib combination showed antagonistic effects in both SKNAS-Rs (3-fold) and SKNAS-RR (1.8-fold) cells (**Supplementary Fig. S5A**), the RMC-6236 plus ribociclib combination moderately decreased IC_50_ values in SKNAS-RR cells (0.36-fold) compared to RMC-6236 monotherapy (**Supplementary Fig. S5B**). Finally, we tested whether SK-N-DZ cells, which harbor a CDK alteration (90), could be re-sensitized in combination with ribociclib. While SK-N-DZ were highly sensitive to ribociclib alone (0.09-fold), the addition of RMC-6236 showed moderate additivity (0.04-fold) compared to RMC-6236 monotherapy (**Supplementary Fig. S5C**). Taken together, these results demonstrate that RMC-6236-resistant cells can be re-sensitized upon combinations with other targeted therapies like TNO155 or ribociclib, and to a larger extent, with avutometinib combinations.

### Efficacy of dual RMC-6236 and avutometinib treatment in neuroblastoma xenografts with acquired RMC-6236 resistance

To determine whether avutometininb can restore sensitivity to RMC-6236 *in vivo*, we monitored tumor growth of ‘RMC6236-resistant’ SKNAS-RR xenografts treated with RMC-6236 (25 mg/kg q.d.), avutometinib (0.3 mg/kg q.d.) or the combination treatment, and monitored drug tolerability, tumor volume and event-free survival. SKNAS-RR xenografts exhibited moderate sensitivity to avutometinib treatment and, surprisingly, retained moderate sensitivity to the RMC-6236 treatment regimen, as evidenced by tumor growth delay (**Fig. 5A)**. However, in contrast with the ‘RMC-6236-sensitive’ SK-N-AS xenografts which showed 100% regression during RMC-6236 treatment (**Fig. 3**), tumor regression was only observed in 1/5 of SKNAS-RR xenografts treated RMC-6236 monotherapy (**Fig. 5B**). Moreover, within 1 week of combination treatment with avutometinib, we observed 5/5 tumor regression that was sustained for over 20 days (**Fig. 5A** and **5B, Supplementary Fig. S6A**). Notably, the combination treatment regimen was optimized at days 20 and 38 to prevent cumulative toxicity, as evidenced by murine weight loss (**Supplementary Fig. S6B**). Finally, event-free survival for the combination-treated group was significantly extended compared to control or monotherapies, with a median survival of 41 days. (**Fig. 5C**).

**Figure 5.**
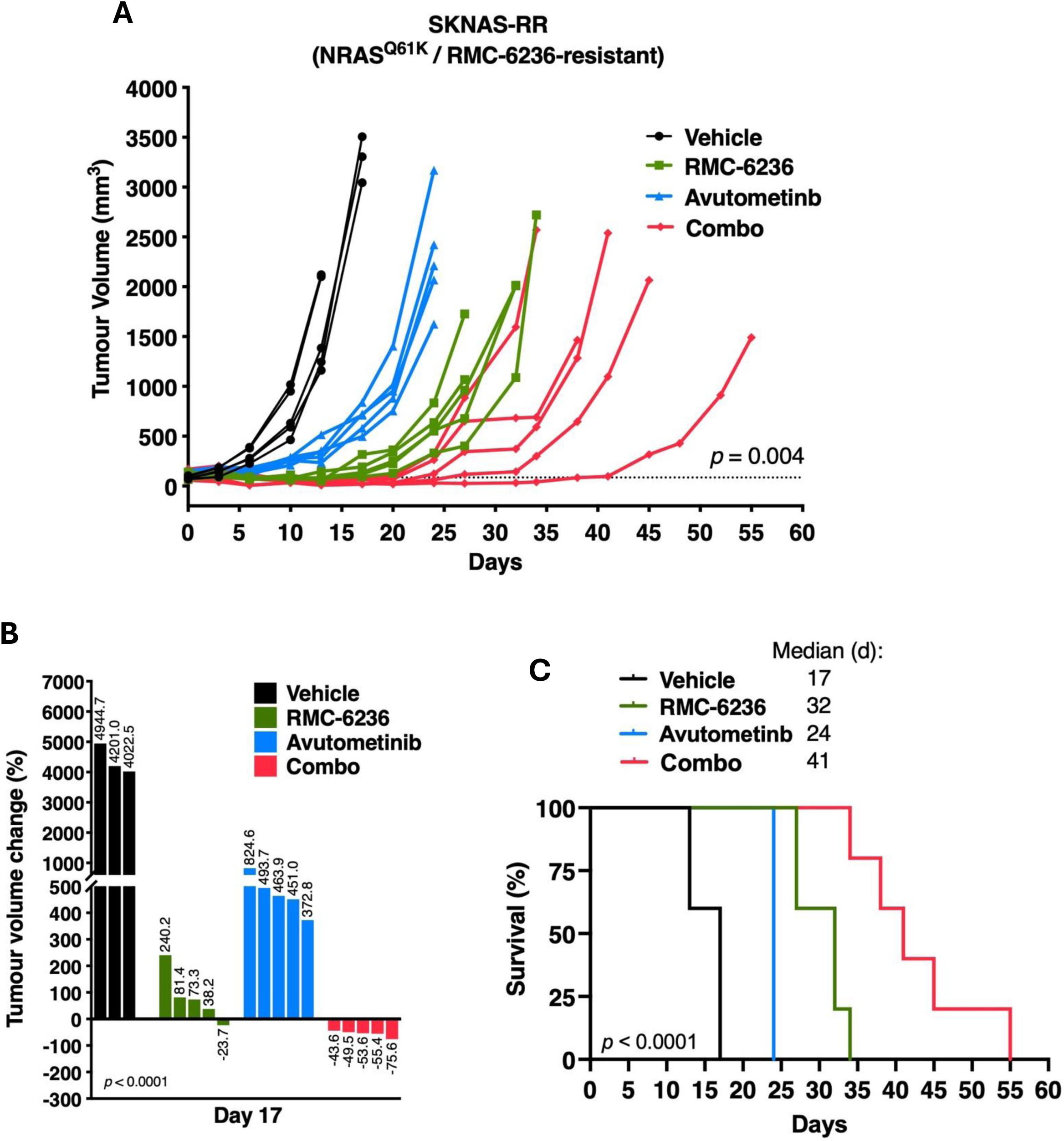
*In vivo* efficacy of RMC-6236 and avutometinib combination in RMC-6236-resistant NB xenografts. **A,** Tumor volumes of SKNAS-RR xenografts (RMC-6236-resistant, NRAS^Q61K^) treated with vehicle (n=5), RMC-6236 (25 mg/kg, q.d., n=5), avutometinib (0.3 mg/kg, q.d., n=5), or combination treatment and monitored until end-point. **B**, Percent tumor change (relative to pre-treatment) following treatment with vehicle, RMC-6236, avutometinib or combination for 17 days. **C**, Kaplan-Meier curve of endpoint-free survival.

### RAS/MAPK, TNFα/NFκB and IL-6/JAK/STAT3 pathway enrichment in RMC-6236 resistance models

To further elucidate molecular mechanisms and specifically changes in pathways associated with RMC-6236 resistance, we next performed transcriptome profiling via RNA sequencing (RNA-seq) comparing gene expression in SKNAS-RR cells, and the more sensitive SKNAS-Rs counterpart. We assessed changes in hallmark signatures and identified 14 pathways that were exclusively and significantly upregulated in SKNAS-RR cells. Among these, SKNAS-RR cells displayed strong enrichment of genes associated in TNFα/NFκB, KRAS, and IL-6/JAK/STAT3 signaling (**Fig. 6A**). Interestingly, although gene set enrichment of epithelial-mesenchymal transition (EMT) was observed (**Supplementary Fig. S7A)**, we did not identify any association with adrenergic (ADRN) or mesenchymal (MES) signatures in SKNAS-RR cells (**Supplementary Fig. S7B)**.

**Figure 6.**
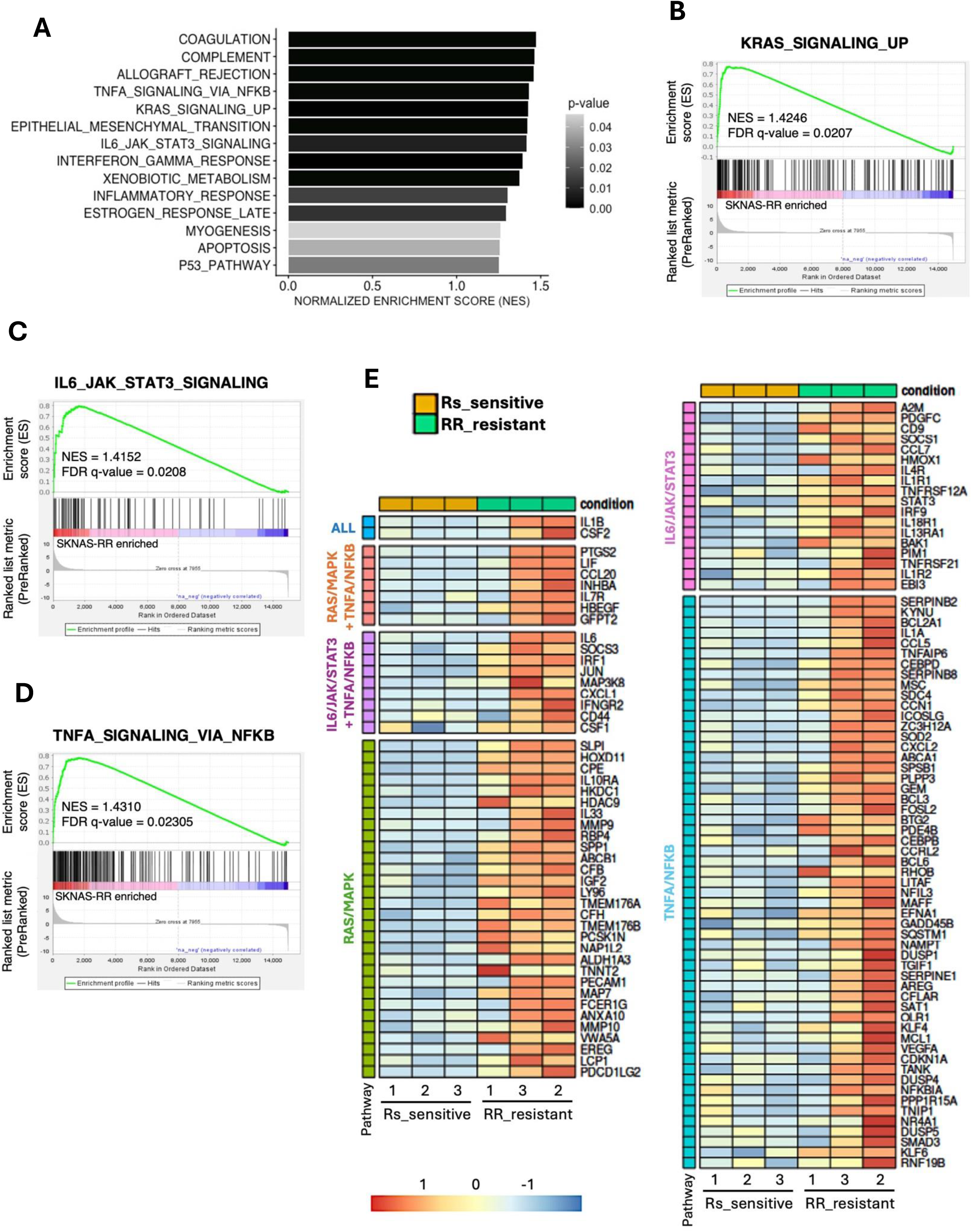
Resistance to RMC-6236 associates with RAS/MAPK, JAK/STAT3 and NFκB pathway enrichment. **A,** Top significantly enriched Hallmark gene sets (MSigDB Database) in SKNAS-RR cell subpopulation following RNA sequencing and displayed according to normalized enrichment scores (NES). Bar color shows statistical significance (NOM p-values < 0.05). **B**-**D**, Gene set enrichment analysis (GSEA) of SKNAS-RR cells showed enrichment of RAS/MAPK (**B)**, TNF⍺/NFκB (**C**), and JAK/STAT3 (**D**) gene-expression signatures. NES and False Discovery Rate (FDR) values are indicated. **E**, Heatmaps of top enriched genes from RAS/MAPK, TNF⍺/NFκB, and JAK/STAT3 signatures and grouped based on pathway overlap: genes shared in all three signatures (‘ALL”, blue), genes common between two pathway signatures (orange and purple), and genes enriched in only one pathway (green, pink and light blue).

Consistent with our previous observations *in vitro* (**Fig. 4C**), GSEA analyses confirmed significant enrichment of the RAS-upregulated gene signature in SKNAS-RR compared to SKNAS-Rs cells (**Fig. 6B**), thus suggesting that combinations with avutometinib are effectively targeting novel vulnerabilities that lead to further activation of RAS/MAPK signaling in our RMC-6236 resistant NB cell model. Furthermore, strong enrichment of IL-6 and JAK/STAT3 pathways (**Fig. 6C**) are in line with previous reports of resistance mechanisms to RMC-6236 (91). Interestingly, SKNAS-RR cells also displayed upregulation of genes involved in TNFα and NFκB signaling, which has been previously shown to contribute to resistance to chemotherapy and other drugs in neuroblastoma (92) (**Fig. 6D**). Finally, to identify other targetable drivers of RMC-6236 resistance we created heatmaps of the top enriched genes from the RAS/MAPK, TNF⍺/NFκB, and JAK/STAT3 signatures and grouped them according to pathway overlap, including genes shared across all three pathways (**Fig. 6E**). Overall, 18 genes were involved in at least two signaling pathways, including *IL1B* and *CSF2* which were common in all three signatures. These genes may represent key mediators of alternative interacting pathways and suggest potential therapeutic targets for combination strategies.

## DISCUSSION

Fewer than half of all high-risk neuroblastoma patients are cured with current chemotherapy, radiation, and immunotherapy treatments. Relapsed and refractory neuroblastoma is even more challenging with fewer than 5% of patients achieving long-term remission. Despite advances in precision oncology and the availability of many new targeted inhibitors and immunotherapies, few drugs are developed for mutations or molecular targets that are only detected in neuroblastoma and/or other pediatric tumors. The development of ALK inhibitors was advanced in large part due to the finding that numerous adult-onset cancers, including subsets of NSCLC, harbor *ALK* rearrangements. Following promising NB efficacy in a Phase II trial (40) the ALK inhibitor lorlatinib is currently being evaluated in a Phase III trial for newly diagnosed neuroblastoma patients with *ALK* activating mutations and/or amplification (NCT03126916). Thus, leveraging promising agents with activity in adult tumors that target genetic alterations that are also detected in neuroblastoma is an important strategy to improve access to novel drugs and potentially reduce the time to first in-child evaluation for NB patients (93).

Increasing data in newly diagnosed and recurrent neuroblastoma suggests that aberrant RAS-MAPK signaling and activation is associated with therapy resistance and poor outcome (33, 35), thus making RAS-MAPK pathway components potential targets for monotherapy or ideally, combination approaches. Following decades of disappointing studies leading to the assumption that RAS was ‘undruggable’, significant progress has been recently made towards successfully targeting active and/or mutation-specific RAS proteins (50, 69, 71, 94–96). Some of the most successful agents include those that specifically target KRAS-G12C variants (97, 98), and although there is evidence for its efficacy in pediatric cancer cell lines, including NB harboring *KRAS-G12C* mutation, these variants only represent a small subset of RAS/MAPK alterations detected in newly diagnosed and recurrent NB. In contrast, RAS(ON) multi-selective inhibitors (e.g. RMC-6236) which target all active GTP-bound RAS isoforms, have been shown to have pre-clinical activity, and efficacy in patients across a wide spectrum of *KRAS*, *HRAS*, and *NRAS* variants, and are of particular interest in tumors that harbor *NRAS* mutations, as there are currently no effective NRAS mutant-specific inhibitors. For instance, the RAS(ON) inhibitor RMC-7977, a preclinical version of RMC-6236, has shown potent antitumor activity via RAS signaling inhibition in *NRAS*-mutant melanomas (99). Additionally, inhibitors like tovorafenib or avutometinib directed against downstream RAS effectors RAF or MEK, have also shown promising results in patients harboring various RAS/MAPK alterations (76, 86). Since relapsed neuroblastoma tumors have a high prevalence of RAS-MAPK pathway alterations, including commonly NRAS mutations, these pan-RAS and MAPK inhibitors are ideal agents to evaluate in our RAS/MAPK-altered neuroblastoma models.

In this study, we assessed the efficacy of tovorafenib, RMC-6236, and avutometinib in neuroblastoma, as these agents are either currently approved for clinical use or under clinical investigation, including several in pediatric age populations (88, NCT04775485, NCT05379985, NCT06104488). While our studies demonstrated that RMC-6236 and avutometinib showed selectivity towards MAPK-altered neuroblastoma cells, tovorafenib treatment did not result in significant cytotoxicity in NB cells harboring *RAS*-, *ALK*– or *NF1* alterations. This was somewhat unsurprising, as previous studies have reported little-to-negligible responses to tovorafenib in other tumors harboring *NRAS* mutations or *NF1* LOF alterations (73, 76). Conversely, selective efficacy of tovorafenib has been demonstrated in tumors bearing *BRAF* mutations or fusions, which are very uncommon in NB cell lines and tumors. Interestingly, our Kelly-LR cells did not respond effectively to tovorafenib despite harboring a BRAF-G466E mutation. This might be due to BRAF-G466 being a class III kinase-impaired mutation that predominantly signals through RAS-dependent CRAF dimerization rather than autonomous BRAF activity, thus reducing its sensitivity, as well as its binding ability, to pan-RAF type II inhibitors such as tovorafenib (100, 101)

In contrast to our tovorafenib findings, results for RMC-6236 support it’s efficacy as a promising therapeutic approach for RAS-MAPK-altered NB, particularly those harboring *RAS*, *NF1* or *ALK* alterations. This is in line with other reports that have showed increased efficacy of RAS(ON) inhibitors in pediatric cancer models harboring rare RAS-G12C alterations, including the LAN-6 neuroblastoma cell line (102). Importantly, RMC-6236 retained cytotoxic activity in the presence of NRAS-Q61K mutation, a common NB variant that we showed conferred resistance to other pathway inhibitors including those targeting SHP2 (67). It’s predecessor, the RAS(ON) inhibitor RMC-7977, showed similar efficacy in *NRAS*-mutant melanomas harboring Q61L or Q61R mutations (99). Additionally, our preclinical studies with RMC-6236 demonstrated tolerability and marked tumor regression in three different RAS/MAPK-mutant NB xenografts. Although our studies unsurprisingly demonstrated tumor regrowth following discontinuation of treatment, there was moderate efficacy following re-treatment. These findings support the selectivity of RMC-6236 for molecularly defined neuroblastoma subsets harboring MAPK alterations, but also highlight the need to identify combination therapies that may prevent or reverse potential resistance to improve long-term responses.

Tumor adaptation through compensatory signaling activation, pathway rewiring, or selection of therapy-resistant subpopulations, can commonly lead to reduced tumor activity and/or durability. A subset of adult patients treated with RAS targeting therapies, including RMC-6236, have been shown at recurrence to acquire DNA alterations predicted to confer resistance including variants in N/H/K RAS, RasGAP, RAF-MEK-ERK and PI3K-AKT-mTOR pathways (99, 103, 104). Similarly, for NB patients several reports have identified emergence in RAS-MAPK alterations during treatment with ALK inhibitors (30, 105, 106). In our studies, RMC-6236 showed limited durability in the absence of continued treatment, as tumor recurrence was observed following treatment discontinuation. Moreover, most recurrent tumors were less sensitive to the same RMC-6236 treatment regimen, suggesting potential development of adaptive resistance. To investigate the mechanisms of RMC-6236 resistance, we established RAS mutant cell line models with acquired resistance to RMC-6236 (SKNAS-RR) and based on previous studies of RAS-targeted agent combinations we evaluated the effect of various inhibitors targeting PARP, CDK4/6 or MAPK components. Notably, avutometinib, TNO155 and ribociclib were effective in combination with RMC-6236 and restored sensitivity in RMC-6236-resistant cells. Importantly, our preclinical studies of RMC-6236 and avutometinib combinations demonstrated increased efficacy and *in vitro* and *in vivo* for tumors with decreased sensitivity to RMC-6236 alone, possibly due in part to hyperactivation of MAPK signaling that we observed in the SKNAS-RR cells. These results provide evidence of potential combination approaches for further study in additional models and patients, especially for patients with multiply relapsed NB who may harbor additional RAS/MAPK activating variants.

Overall, our findings support RMC-6236 as a promising therapeutic strategy for neuroblastoma, particularly those harboring *RAS*, *NF1* or *ALK* alterations, or MAPK pathway-associated resistance. Our initial studies exploring gene expression differences in RMC-6236-resistant cell lines suggest that mechanisms of resistance may include activation of additional RAS/MAPK signaling and non-MAPK pathways including JAK-STAT and NFκB. These findings and ongoing whole genome studies will provide insight into resistance mechanisms that might be targets for combination therapies with RMC-6236. However, the current findings together with results in adult human trials for RMC-6236 (daraxonrasib) monotherapy and with other agents, support prioritizing further studies for the subset of NB patients with alterations in this pathway. Our results also support consideration of avutometinib combination with RMC-6236 in NB. Although these drugs have not been evaluated together in adult clinical trials there is additional pre-clinical data demonstrating efficacy for the KRAS-G12D targeted agent MRTX1133 combination with avutometinib in PDAC pre-clinical models (79). Further studies of the mechanisms of resistance in cell and animal models as well as ongoing monitoring of evolution of RAS-MAPK variants in the ctDNA from adult patients treated with RMC-6236 will provide additional important data to form the basis of rational RMC-6236 combination therapies in patients with NB and other tumors with activated RAS signaling.

## ACKNOWLEDGMENTS

This project was funded by the Canadian Institutes of Health Research (CIHR) Operating Grant (PJT-162228 to M.S. Irwin and PJT-166005 to M. Ohh), SickKids Neuroblastoma Research Funds including James Fund, Curtis Chow Memorial Fund, Lilah’s Fund, and Sebastian’s Superheroes (M.S. Irwin).

## Disclosure of Potential Conflicts of Interest

The authors declare no potential conflicts of interest.

## Financial Support

This project was funded by the Canadian Institutes of Health Research and James Fund for Neuroblastoma Research.

## SUPPLEMENTARY FIGURE LEGENDS

**Supplementary Figure S1.**
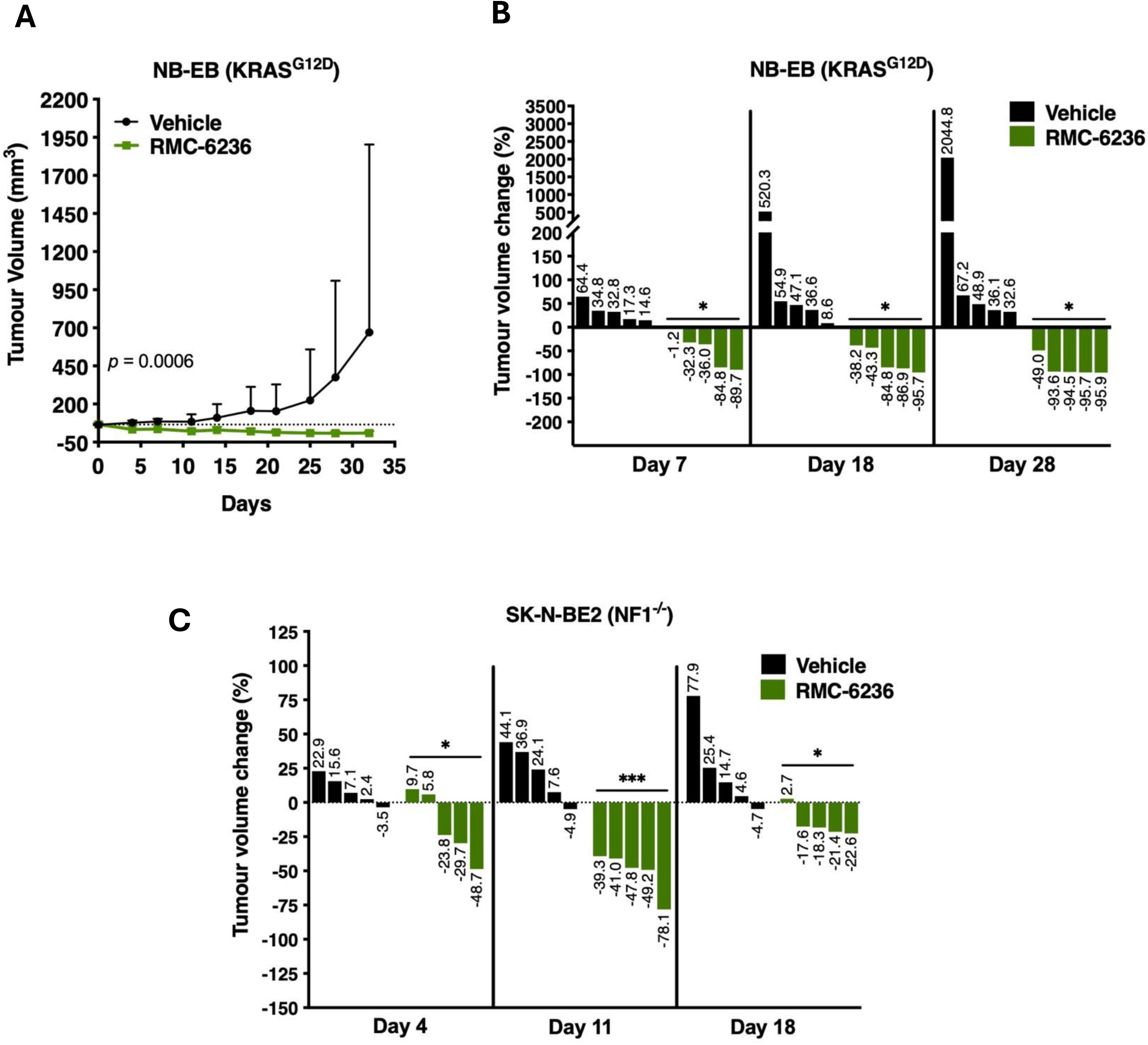
*In vivo* efficacy of RMC-6236 in *RAS-* and *NF1-*altered xenografts. A-B, Tumor volumes (**A**) and percent tumor volume change (relative to pre-treatment) (**B**) of NB-EB xenografts treated with vehicle (n=5) or RMC-6236 (25 mg/kg, q.d., n=5) for the indicated days. **C**, Percent tumor volume change of SK-N-BE2 xenografts following treatment with vehicle (n=5) or RMC-6236 (25 mg/kg, q.d., n=5) for the indicated days.

**Supplementary Figure S2.**
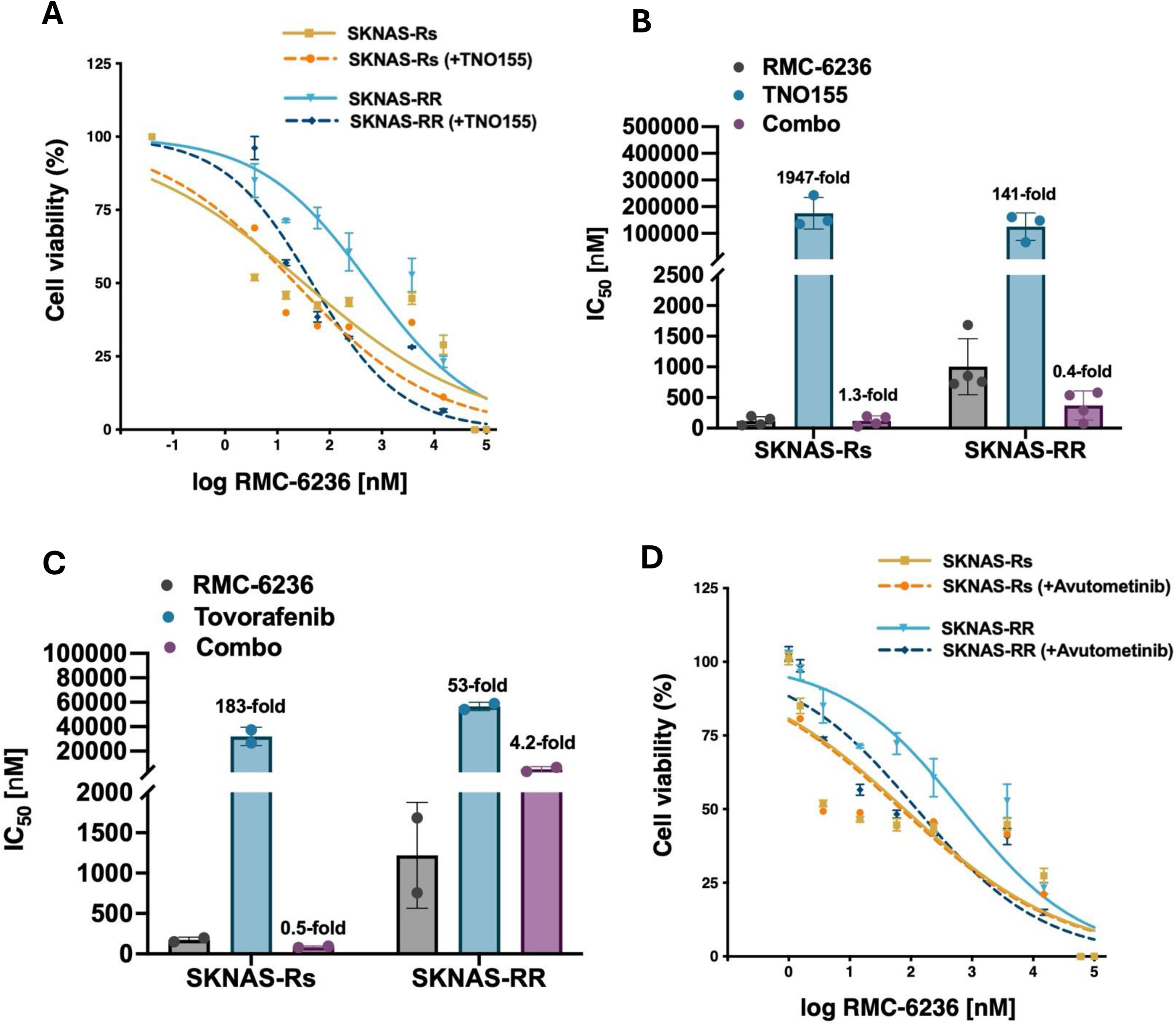
Assessment of RMC-6236 combinations with MAPK inhibitors in SKNAS-RR cells. **A**, Cell viability of SKNAS-Rs and SKNAS-RR cells treated with wide-range concentrations of RMC-6236 plus anchored TNO155 [5 μM] for 120 hours. **B**-**C**, Average IC_50_ and fold change (relative to RMC-6236 alone) of SKNAS-Rs and –RR cells treated with RMC-6236, TNO155 or Combo (**B**), or RMC-6236, tovorafenib or Combo (**C**) for 120 hours. **D**, Cell viability of SKNAS-Rs and –RR cells treated with wide-range concentrations of RMC-6236 plus anchored avutometinib [0.2 μM] for 120 hours.

**Supplementary Figure S3.**
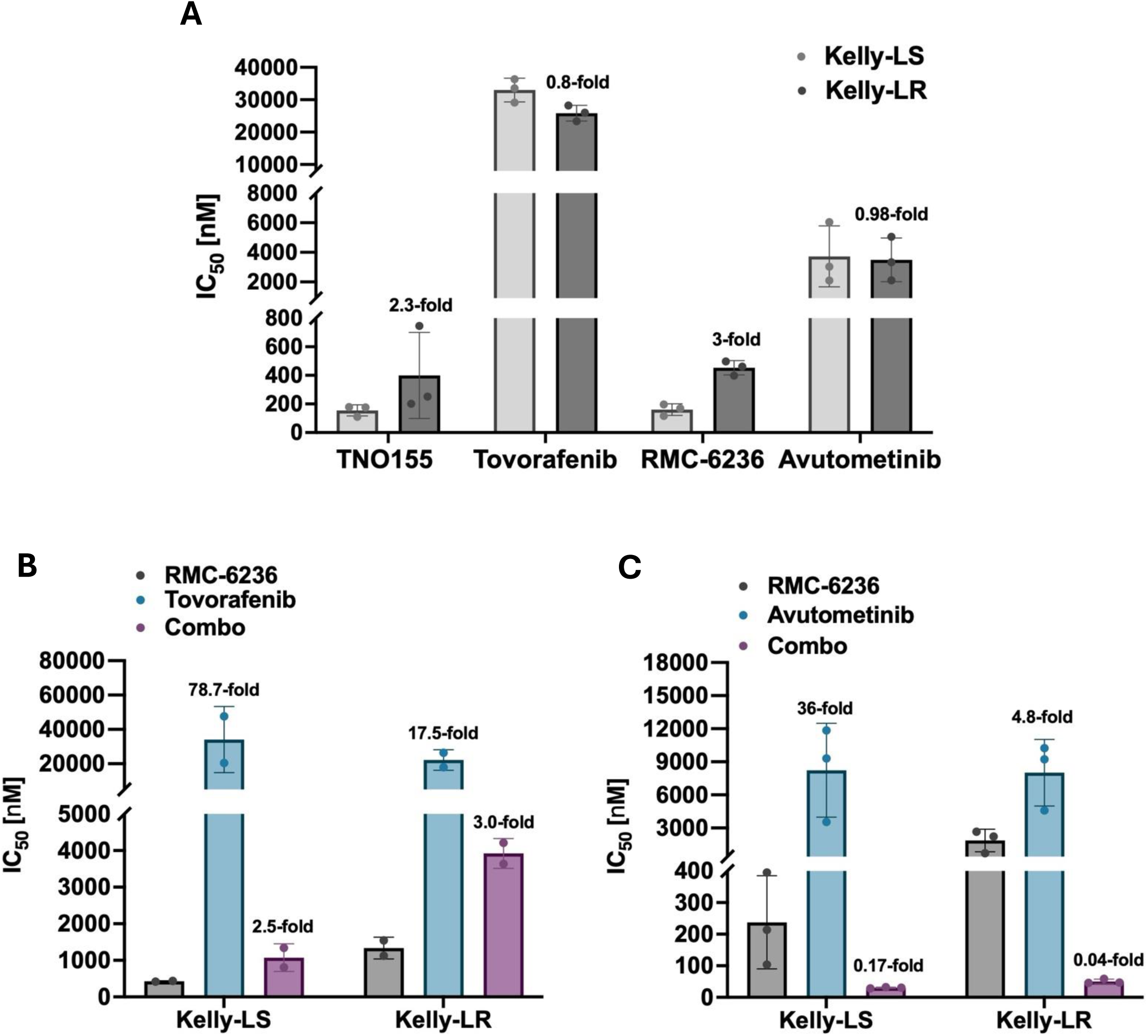
*BRAF*-mutant Kelly-LR (lorlatinib-resistant) cells are sensitive to avutometinib and RMC-6236 combination treatment. **A**, Average IC_50_ of lorlatinib-sensitive (Kelly-LS) and lorlatinib-resistant (Kelly-LR) cells treated with TNO155, tovorafenib, RMC-6236, or avutometinib for 120 hours. IC_50_ fold change relative to Kelly-LS cells. **B**-**C**, Average IC_50_ of Kelly-LS and Kelly-LR cells treated with RMC-6236, tovorafenib or Combo (**B**) or RMC-6236, avutometinib or Combo (**C**) for 120 hours. IC_50_ fold change relative to RMC-6236 treatment alone.

**Supplementary Figure S4.**
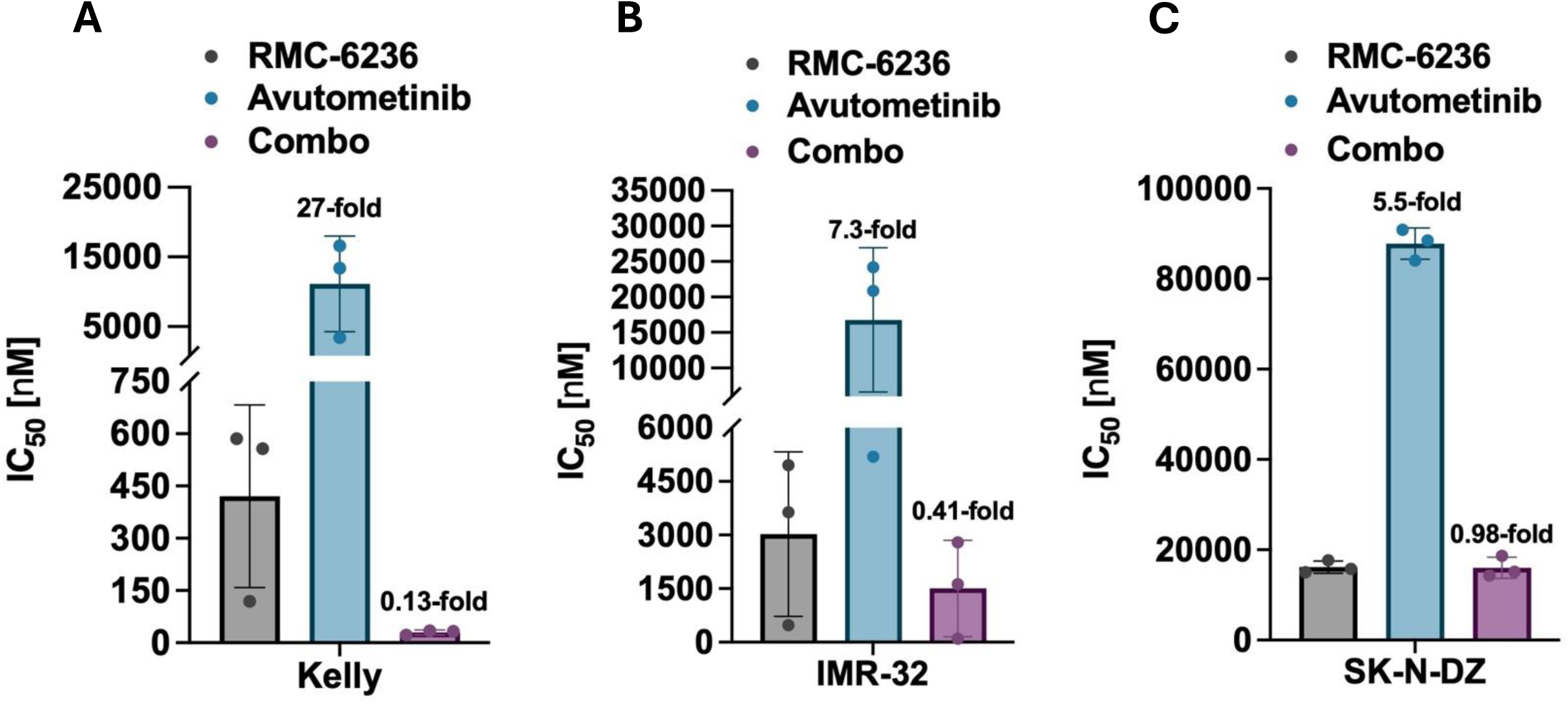
NB cell lines with intrinsic resistance to RMC-6236 are sensitive to avutometinib plus RMC-6236 treatment. A-C, Average IC_50_ and fold change (relative to RMC-6236 alone) of Kelly (**A**), IMR-32 (**B**) or SK-N-DZ (**C**) cells treated with RMC-6236, avutometinib or Combo for 120 hours. IC_50_ fold change relative to RMC-6236 treatment alone.

**Supplementary Figure S5.**
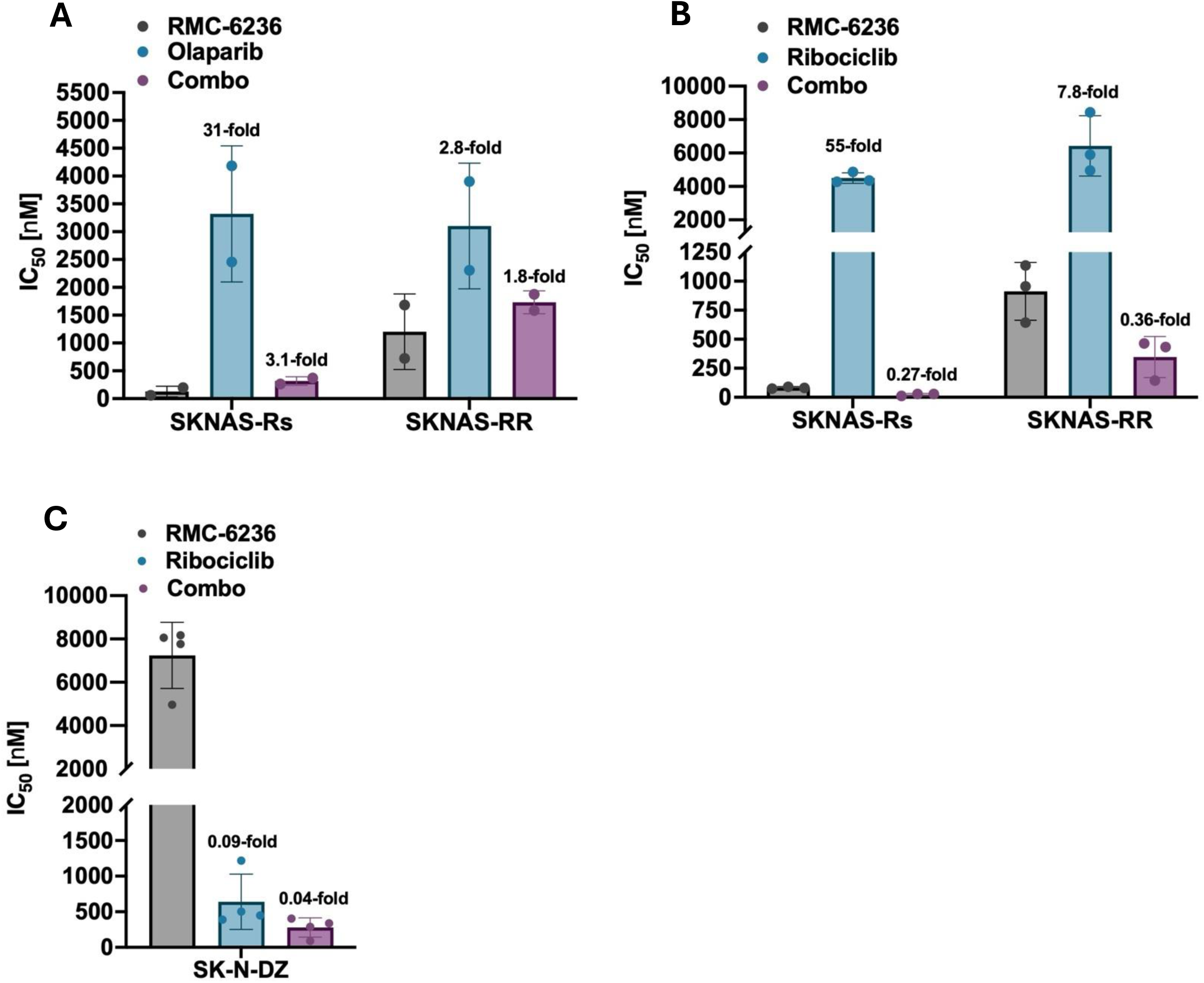
Assessment of RMC-62636 combinations with PARP and CDK4/6 inhibitors. **A**, Average IC_50_ of SKNAS-Rs and SKNAS-RR cells treated with RMC-6236, olaparib or Combo (**A**) for 120 hours. **B**-**C,** Average IC_50_ of SKNAS-Rs and –RR cells (**B**) or SK-N-DZ cells (**C**) treated with RMC-6236, ribociclib or Combo for 120 hours. IC_50_ fold change relative to RMC-6236 alone.

**Supplementary Figure S6.**
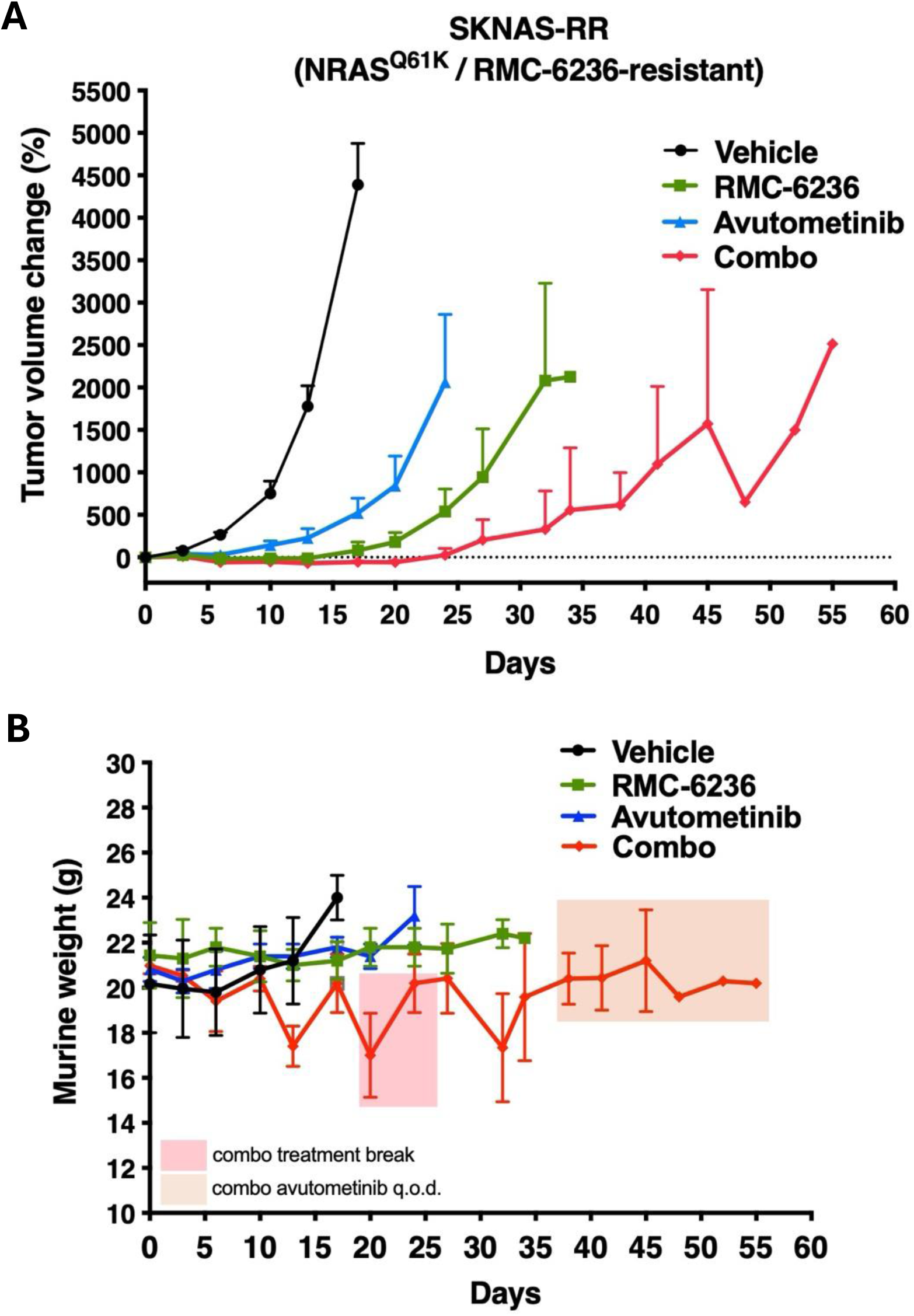
Efficacy of RMC-6236 plus avutometinib in RMC-6236-resistant NB xenografts. **A,** Percent tumor volumes change of SKNAS-RR xenografts (RMC-6236-resistant, RR) treated with vehicle (n=5), RMC-6236 (25 mg/kg, q.d., n=5), avutometinib (0.3 mg/kg, q.d., n=5), or combination treatment and monitored until end-point. **B,** Average murine weight of SKNAS-RR xenografts. Combo treatment break occurred from days 20 to 26 after observing weight loss. At day 38 the combination treatment group received RMC-6236 (25 mg/kg, q.d.) plus avutometinib (0.3 mg/kg, every other day q.o.d.). Neither monotherapy treatment had a treatment regimen change throughout the study.

**Supplementary Figure S7.**
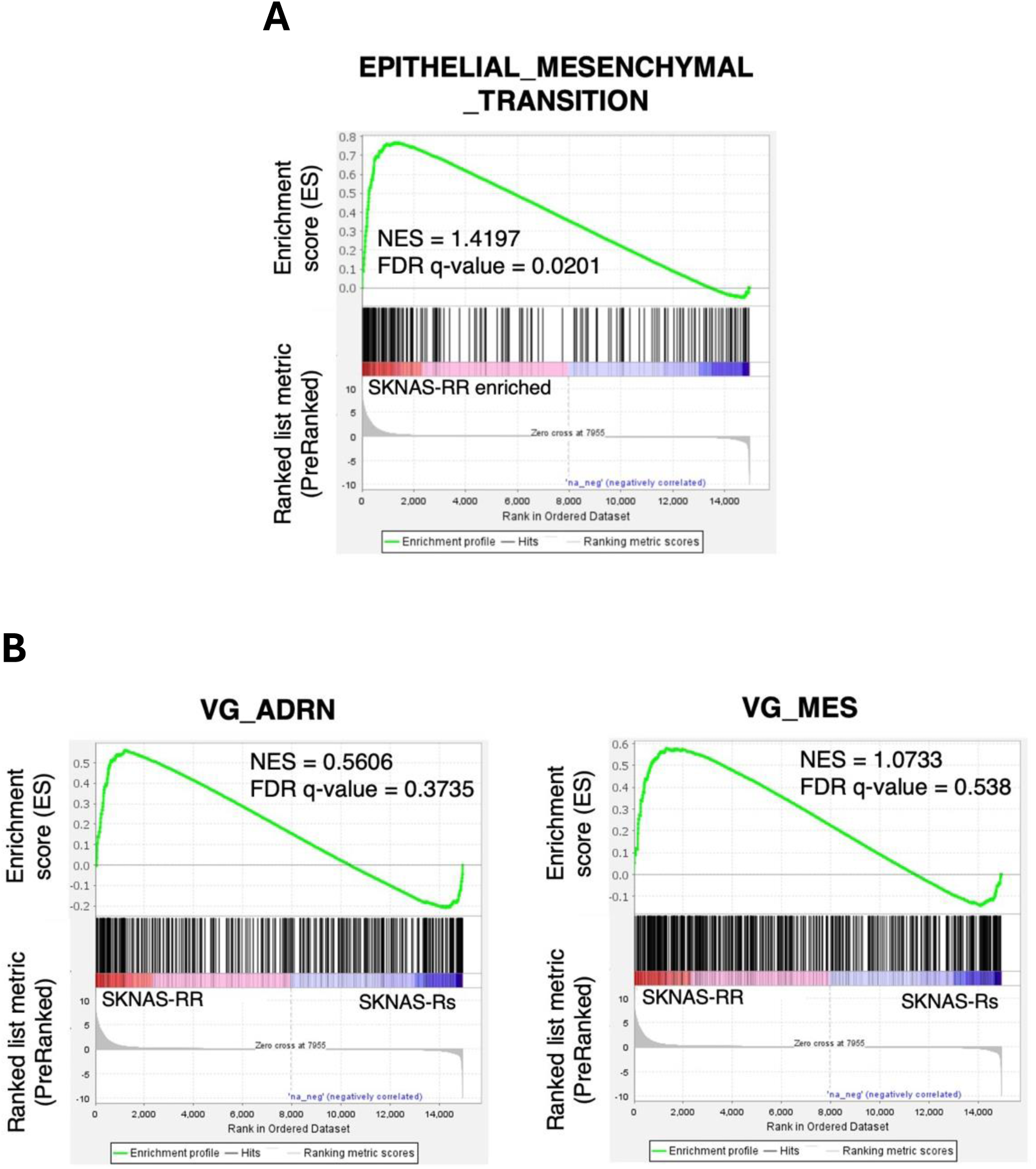
Enrichment of EMT pathway in SKNAS-RR cells. **A**-**B**, Gene set enrichment analysis (GSEA) of SKNAS-RR cells indicates enrichment of epithelial to mesenchymal transition (EMT) gene-expression signatures (**A**), but not of adrenergic (ADRN) or mesenchymal (MES) signatures (**B**). Normalized enrichment scores (NES) and False Discovery Rate (FDR) values are indicated.

## Notes

### Competing Interest Statement

The authors have declared no competing interest.

